# Optogenetic analysis of Ca^++^ transients in *Caenorhabditis elegans* muscle cells during forward and reverse locomotion

**DOI:** 10.1101/2020.12.02.408088

**Authors:** Jacob R. Manjarrez, Magera Shaw, Roger Mailler

## Abstract

Understanding how an organism generates movement is an important step toward determining how a system of neurons produces behavior. With only 95 body wall muscles and 302 neurons, *Caenorhabditis elegans* is an attractive model organism to use in uncovering the connection between neural circuitry and movement. This study provides a comprehensive examination of the muscle cell activity used by *C. elegans* during both forward and reverse locomotion. By tracking freely moving worms that express genetically encoded calcium indicators in their muscle cells, we directly measure the patterns of activity that occur during movement. We then analyzed these patterns using a variety of signal processing and statistical techniques. Although our results agree with many previous findings, we also discovered there is significantly different mean Ca^++^ levels in many of the muscle cells during forward and reverse locomotion and, when considered independently, the dorsal and ventral muscle activation waves exhibit classical neuromechanical phase lag (NPL).

## INTRODUCTION

One of the primary goals of neuroscience is to uncover the relationship between neural organization and behavior. A key contribution toward achieving this goal occurred with the publication of the first nearly complete connectome of the nematode *Caenorhabditis elegans* (White, et al., 1986). Since that time, researchers have simultaneously worked to fill in the missing details of the neural map (Varshney, et al., 2011; Cook, et al., 2019) and relate its structure to behaviors such as chemotaxis (Chalasani, et al., 2007), touch response (Goodman, 2006), and locomotion (Karbowski, et al., 2008; Boyle, et al., 2007; Wen, et al., 2012; Tolstenkov, et al., 2018; Fouad, et al., 2018).

Traditionally, the activity of the 95 body wall muscles (BWM) that generate locomotive force has been inferred by measuring changes in body posture as the worm moves (Baek, et al., 2002; Cronin, et al., 2005). This has led researchers to hypothesize that *C. elegans* undulating locomotion is generated through alternating dorsal/ventral muscle waves that propagate at the same rate as the body wave from head to tail during forward locomotion and from tail to head during reverse locomotion (Niebur & Erdos, 1991; Boyle, et al., 2007; Haspel, et al., 2011; Wen, et al., 2012). Furthermore, the ventral nerve cord A- and B-type neurons have been implicated as the primary muscle drivers, while the D- and AS-type neurons act to prevent co-activation of opposing dorsal/ventral muscles (Haspel, et al., 2011; Wen, et al., 2012; Tolstenkov, et al., 2018).

In this study, we investigate the behavior of the BWMs during unrestrained forward and reverse locomotion by measuring their Ca^++^ activity using the genetically encoded calcium indicator GCamp3. The data collected during these experiments are then subjected to a battery of statistical and signal analysis techniques. The results of the analysis agree with many previously published findings, but also reveal new important discoveries about the muscle waves generated by *C. elegans*. For example, our results support the conclusions that the head acts nearly independently of the body (Alkema, et al., 2005; Fouad, et al., 2018), that there is a linear drop-off in muscle activation during wave propagation (Xu, et al., 2018), that the wave frequencies of the muscle cells matched those of the bend angles (Karbowski, et al., 2008), and in agreement with the work of Butler et al. (Butler, et al., 2015), that the neuromuscular wave travels faster, and with a fixed phase offset, than the body bend wave.

However, our results also show that dorsal/ventral muscle activation is only truly anti-phasic in the mid-body and shows a larger distribution in other parts of the body than previously reported (Pierce-Shimomura, et al., 2008; Butler, et al., 2015). The results further show that, when considered individually, the ventral and dorsal neuromuscular waves propagate down the body at different rates from each other and exhibit a prototypical neuromechanical phase lag (NPL) like those found in many anguilliform swimmers (McMillen, et al., 2008; Tytell, et al., 2010). Our analysis shows that the muscle waves are initially anti-phasic but realign themselves as they progress from head to tail during forward locomotion, with the exact opposite occurring during reversals.

Finally, when comparing forward to reverse locomotion, our results show that there are significant differences in the pattern of calcium levels throughout the BWMs. During forward locomotion, for example, both dorsal and ventral muscle groups exhibit significantly higher mean anterior calcium levels, which linearly taper off until they become significantly lower in the posterior (Xu, et al., 2018). The exact opposite phenomena occur during reverse locomotion.

## RESULTS

### Muscle cell activity decreases linearly as the locomotion wave propagates from anterior to posterior during forward locomotion and posterior to anterior during reverse locomotion

Raw intensity measurements (Figures 1B & C) show high levels of calcium activity in the midbody of the worm during forward locomotion and near the tail during reverse locomotion. However, when normalized (see Methods) against measurements taken of worms exposed to 4AP (Figure 1A), the normalized mean values (Figures 1D & E) reveal that muscle cells rarely exceed 50% of the maximum levels. In addition, they show that the dorsal and ventral sides have equal mean levels of Ca^++^ intensity (α=0.01). During forward movement, activity in the head is higher than the rest of the body. The average normalize intensity for forward locomotion is 47.3 ± 9.7% in the head and falls off to 17.3 ± 5.7% in the tail. This represents a falloff of over 63% over the length of the body and is well fit by the linear equation *foract(x) = −0.0094x + 0.469* with an R^2^ = 0.83. These results show that during forward locomotion, the dorsal and ventral mean Ca^++^ levels are highest in the head and decrease distally, and that the mean levels are simultaneously elevated on both the dorsal and ventral sides.

**Figure 1:**
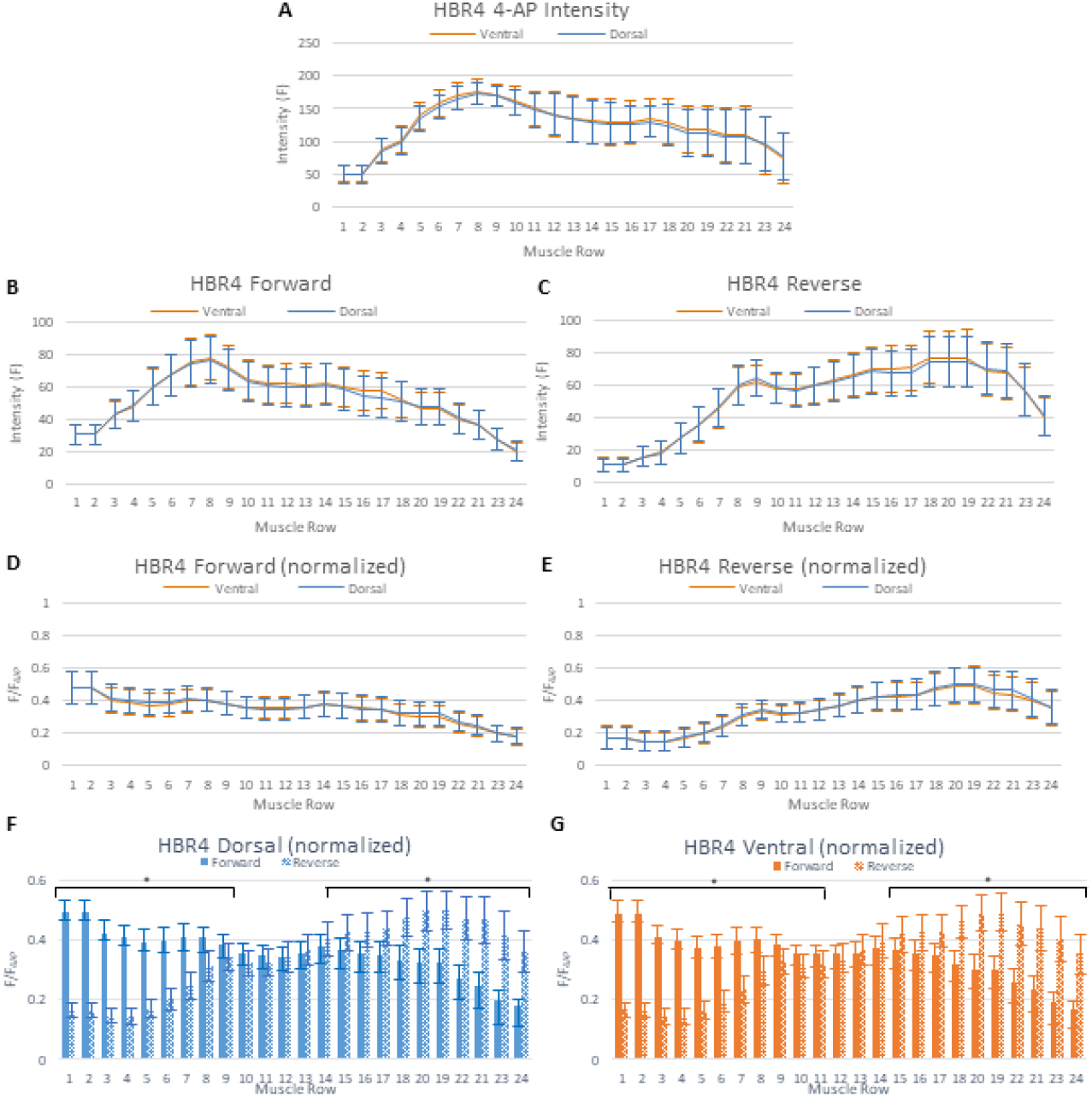
The normalized average GFP intensity along the AP axis during forward and reverse locomotion. Average and normalized fluorescence intensities recorded from HBR4 worms. Solid lines indicate the average and bars indicate the standard deviation. A) The average GFP intensities from the dorsal and ventral muscles for worms exposed to 500mM 4-AP. Averages recorded from this data were used to normalize GFP intensities. B) The average (not normalized) GFP intensities for the dorsal and ventral muscles along the AP axis during forward locomotion. C) The average (not normalized) GFP intensities for the dorsal and ventral muscles along the AP axis during reverse locomotion. D) The average normalized GFP intensities recorded from HBR4 worms during forward locomotion. The normalized intensities drop off approximately linearly from head to tail. E) The average normalized GFP intensities recorded from HBR4 worms during reverse locomotion. The normalized intensities drop off approximately linearly from tail to head. E) The average normalized Ca++ intensities for the dorsal muscles during forward and reverse locomotion. The asterisk (*) indicates a significant difference (α = 0.01) between forward and reverse intensities. F) The average normalized Ca++ intensities for the ventral muscles during forward and reverse locomotion. The asterisk (*) indicates a significant difference (α = 0.01) between forward and reverse intensities.

The normalized standard deviations of these signals also decrease linearly (decreasing ~41%) over the length of the worm. Since the amplitude of a sine wave is proportional to its standard deviation and can be calculated using the equation (Smith, 1999),

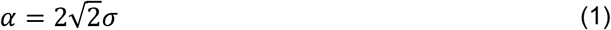

our results show that the peak-to-peak amplitude of the dorsal and ventral waves are highest in the head and deceases along the length of the body. These results agree with the findings of Xu et al. who presented a linear model that predicts a deteriorated undulatory wave in the presence of AVB-B gap junctions (Xu, et al., 2018).

During reverse locomotion (Figure 1E), the pattern is nearly perfectly reversed. Normalized Ca^++^ levels in the tail are higher than the rest of the body, as are the standard deviations. In the tail, the average normalized intensity is as high at 49.5 ± 11.2% at muscle row 19 while in the head it falls to just 17.1 ± 7.1%. This represents a 65% falloff in the mean intensity and a 37% decrease in the standard deviation over the length of the worm. The linear equation *revact(x) = 0.0151x + 0.153* with an R^2^ = 0.81 fits the falloff in mean levels well.

When comparing the amplitudes of the Ca^++^ signals in the ventral and dorsal muscles, the results indicate that there is a significant (α=0.01) difference between their activity during forward and reverse locomotion. As Figure 1F & G show, during forward locomotion, dorsal and ventral muscle rows 1-10 show significantly higher and muscle rows 15-24 show significantly lower mean normalized Ca^++^ intensity levels when compared to reverse. During reverse locomotion, the opposite is true with muscle rows 1-10 exhibiting significantly lower and muscle rows 15-24 exhibiting significantly higher mean normalized Ca^++^ levels on both the dorsal and ventral sides.

### The head acts independently of the rest of the body

During forward locomotion, the muscle cells in the head (rows 1-6) have highly correlated Ca^++^ activity, indicating that they act as a group to control the head position (Figures 2A & B). However, there is an abrupt shift in the patterns between muscle rows 4 and 6 where the ventral and dorsal sides change from phasic to anti-phasic behavior. At muscle row 5, autocorrelation analysis shows alternating patterns of phasic and anti-phasic behavior, which can be seen in the large standard deviations posterior of the nerve ring in Figure 2A. These results appear to indicate that two different neural circuits, which overlap near the nerve ring, are primarily responsible for separate control of the head and body during forward locomotion. These results agree with experiments conducted by Fouad et al. who demonstrated that the head can effectively be disconnected from the rest of the body with minimal impact to the generation and propagation of locomotion waves (Fouad, et al., 2018).

**Figure 2:**
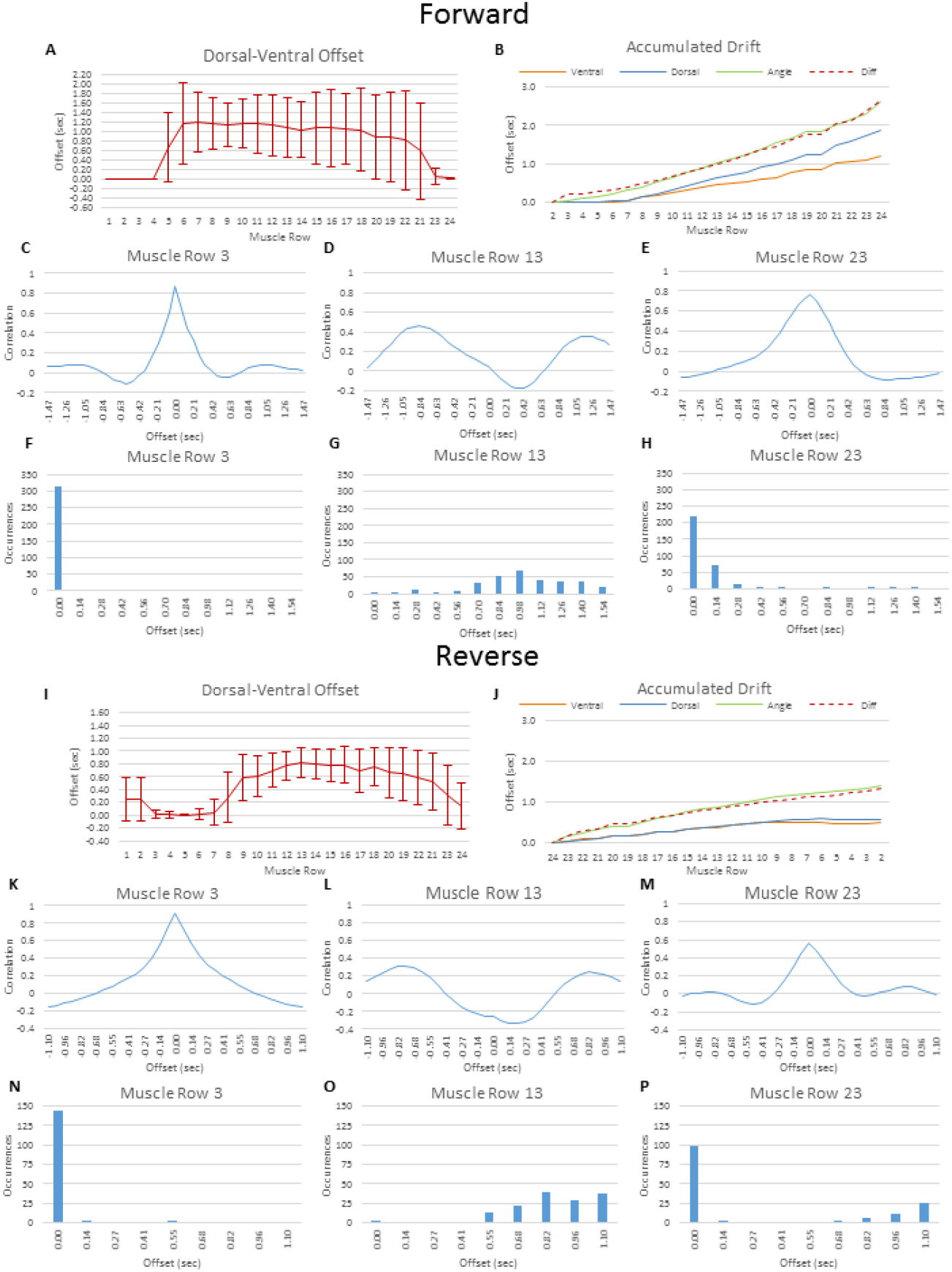
Muscle activation in the head and tail are in phase while activation in the midbody is anti-phasic. The ventral side wave also travels faster than the dorsal side wave. A) Offset between the signals in the dorsal and ventral muscle groups during forward locomotion. Solid line indicates the average offset, while bars indicate the standard deviation. B) The accumulated offset as the wave travels down the AP axis. Solid orange, blue, and green lines are the average accumulated offset for ventral signal, dorsal signal, and bend angle, respectively. The dashed red line shows the difference in activation. C-E) Average cross correlation of the dorsal and ventral muscle activations at the 3^rd^, 13^th^ and 23^rd^ muscle rows during forward locomotion. F-H) Histogram of the maximum point of dorsal/ventral cross correlation recorded from the 3^rd^, 13^th^ and 23^rd^ muscle rows. I) Offsets between the signals in the dorsal and ventral muscle groups during reverse locomotion. J) Accumulated drift from tail to head during reverse locomotion. K-M) Average cross correlation of the dorsal and ventral muscle activations at the 3^rd^, 13^th^ and 23^rd^ muscle rows during reverse locomotion. N-P) Histogram of the maximum point of dorsal/ventral cross correlation recorded from the 3^rd^, 13^th^ and 23^rd^ muscle rows.

During reverse locomotion, head activation is minimal, but muscle cell activity remains largely correlated. Figures 2I & J show very small temporal offsets between the ventral and dorsal sides as well as between subsequent rows of muscles. A pattern transition like the one seen during forward locomotion occurs between muscle rows 9 and 7 with muscle row 8 showing alternating phasic and off-phasic activity. These results support the conjecture that the head is controlled using a separate neural circuit from the rest of the body and previous findings that show that head activity is suppressed during reverse locomotion (Alkema, et al., 2005).

### Muscle activation patterns contain both phasic and anti-phasic components

Within the head of the worm, dorsal and ventral side intensities are directly in phase with one another. Figure 2A shows that in muscle rows 1-4, the dorsal and ventral sides have no offset in signal when evaluated using autocorrelation. Figure 2C shows the results of the average autocorrelation of the dorsal and ventral muscles in row 3. The point of maximum correlation has a 0 second offset. Additionally, in each of the 315 fragments analyzed (Figure 2F), the signals are in perfect synchronization.

On average, within the mid-body of the worm, the dorsal and ventral sides are out of phase (Figure 2A). However, the non-symmetric cross correlation graph in Figure 2D shows that the activations are not completely anti-phasic. Figure 2G also shows that, unlike the muscles in the head, the offset between the dorsal and ventral sides drift considerably in the mid-body. Among the fragments evaluated for muscle row 13, the activation offset has a mean value of 0.96 ± 0.32 seconds. Given the characteristic frequency of 0.44Hz observed in the muscle cells, the expected dorsal and ventral offset should be 1.14 seconds. While an offset of 1.14 seconds occurs near the nerve ring, more posterior the offset decreases.

At the tail of the worm, the Ca^++^ waves are in phase once again (Figure 2A). This is confirmed in Figures 2E and 2H, which show that the average maximum cross correlation occurs at a 0 second offset and that this happens in nearly all the video fragments evaluated.

Figure 2B shows why the muscle Ca^++^ patterns are in phase, then out of phase, then drift back into phase once again. In the head, the offset between muscle rows 1-5 is nearly 0, meaning they occur at the same time. However, posterior of the nerve ring, the dorsal and ventral muscle activation waves travel down the body at different speeds. The ventral wave takes an average of 1.18 seconds to travel down the body while the dorsal wave takes approximately 59% longer or an average of 1.88 seconds. However, when the two waves are subtracted from one another, the resulting combined signal has an average accumulated offset of 2.66 seconds, which closely approximates the average accumulated offset for the body wave of 2.58 seconds.

In reverse locomotion (Figures 2I-P), the same pattern holds although the accumulated drift between the dorsal and ventral sides is less pronounced (Figure 2J). In the head and tail of the worm, the activation of the dorsal and ventral muscles is nearly perfectly in phase while in the mid-body, they become out of phase (Figures 2K-P). Overall, the reverse wave is nearly twice as fast as the forward wave taking only 1.34 versus 2.58 seconds. Like the forward wave, the dorsal and ventral activation waves are twice as fast as the body wave and the difference in the ventral and dorsal activation is also most closely associated with overall body wave propagation.

### Body posture is highly correlated with the difference in intensity between ventral and dorsal muscle activation. Bend speeds are weakly correlated with muscle activity

During forward locomotion, body bending (Figure 3A) deviates from center nearly equally in the dorsal and ventral directions. Overall, bend angle is most highly correlated with the difference in the dorsal-ventral Ca^++^ levels except in the first two muscle rows of the head (Figure 3C). Posterior of the nerve ring, dorsal and ventral muscle Ca^++^ levels are ~50% correlated with bend angle, however as the wave propagates further back, the correlation with ventral activation decreases. Additionally, Fourier analysis (Figures 3E-G) shows that the dominant frequency components (0.11 and 0.44Hz) for dorsal and ventral activation are nearly equal. However, bending and the difference in Ca^++^ level frequencies are most highly correlated throughout the worm’s body with a single dominant frequency of 0.44Hz. These results agree with the findings of Karbowski et al. who reported component bend angle frequencies in the range of 0.45-0.48 Hz (Karbowski, et al., 2008).

**Figure 3:**
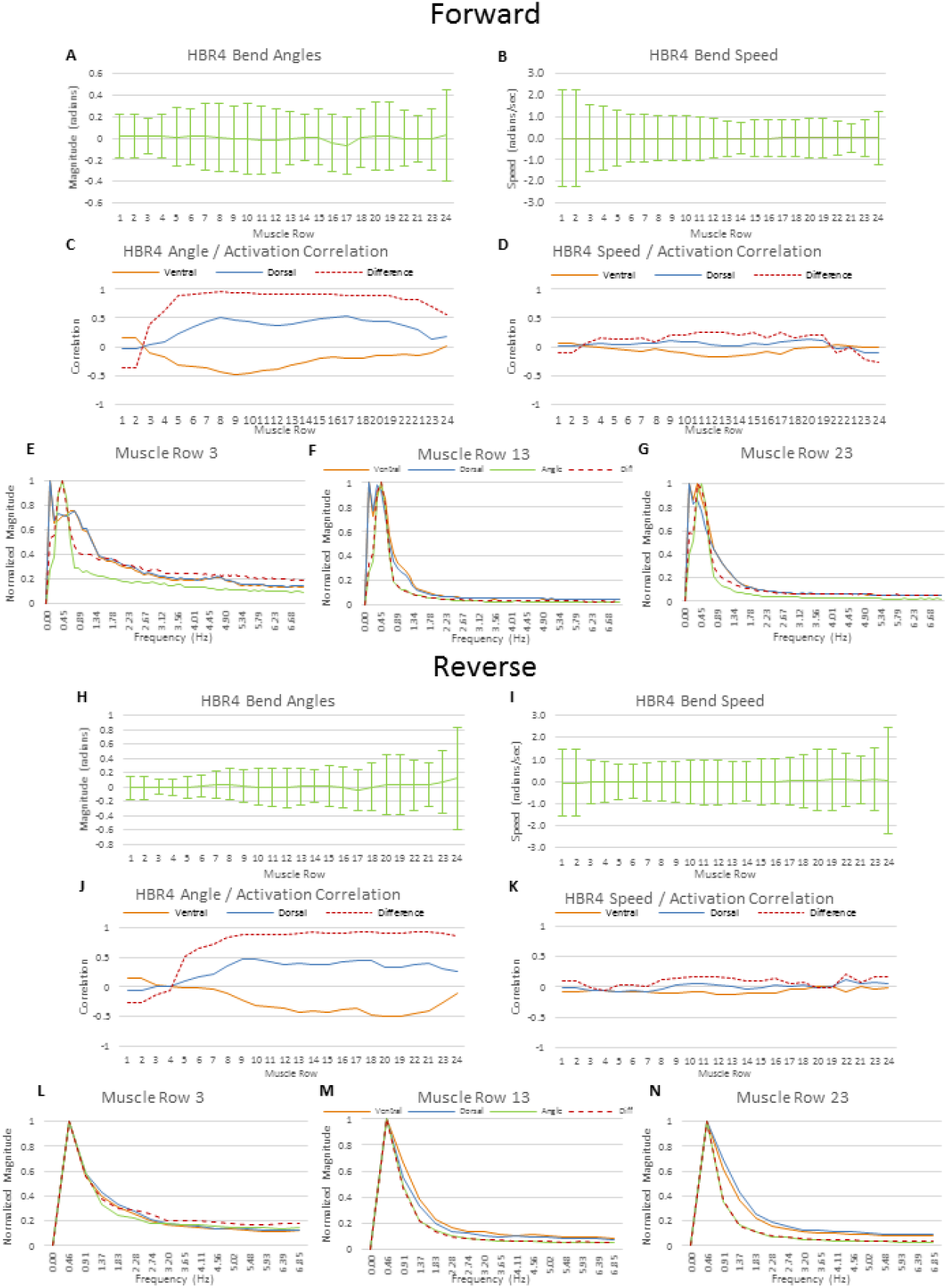
Correlation between body posture, muscle speed, and muscle activation during forward and reverse locomotion. A) Average bend angles of the body of an HBR4 worm during forward locomotion. Solid lines indicate the average and bars indicate the standard deviation. B) Average bend speed of the body of an HBR4 worm during forward locomotion. Solid lines indicate the average and bars indicate the standard deviation. C) Correlation of bend angle with the muscle activation during forward locomotion. Dorsal and ventral correlations are indicated with solid lines, where the difference in activation (D-V) is indicated as a dashed line. D) Correlation of the bend speed with muscle activation during forward locomotion. Dorsal and ventral correlations are indicated with solid lines, where the difference in activation (D-V) is indicated as a dashed line. E-G) Frequency analysis of bend angle, muscle activation, and difference in muscle activation at muscle rows 3, 13, and 23. H) Bend angles of the body of HBR4 during reverse locomotion I) Bend speeds of the body of HBR4 during reverse locomotion. J) Correlation of bend angle with muscle activation during reverse locomotion. K) Correlation of bend angle speed and muscle activation. L-N) Frequency analysis of the bend angle, muscle activation, and difference in muscle activation during reverse locomotion.

In reverse locomotion, the results are similar. Bend angles tend to be the greatest near the tail of the worm (Figure 3H) and remain most highly correlated with the difference in dorsal-ventral Ca^++^ levels (Figure 3J). Frequency analysis (Figure 3L-N) shows that bending frequency is most closely associated with the frequency of the difference between dorsal and ventral Ca^++^ levels.

During forward locomotion, bend angles (Figure 3B) change most quickly in the head and then at nearly equal speeds throughout the rest of the body. During reverse locomotion, bend speeds (Figure 3I) are greatest in the tail and at the tip of the head. In both forward and reverse locomotion, bend speeds show little correlation with muscle Ca^++^ activity (Figure 3D & K).

### Dorsal and ventral muscle activation wave combine to create the overall activation pattern

From a mathematical perspective, we can describe the activity of a muscle cell at time *t* as sine wave with a wavelength (λ), amplitude (*α*), and phase (*ϕ*) using the formula *w*(*t*) = *α* sin(*λt* + *ϕ*). Since the dorsal and ventral muscles work in opposition to one another, the combined activity of these cell can be calculated by subtracting one from the other. In other words, if we use the function *d*(*t*) = *α_d_* sin(*λ_d_t* + *ϕ_d_*) to describe the activity of the dorsal side and *v*(*t*) = *α_v_* sin(*λ_r_t* + *ϕ_v_*) for the ventral side, the resultant activity for the muscle row becomes *r*(*t*) = *d*(*t*) — *v*(*t*). Substituting in we get

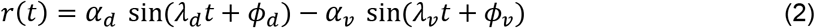

Based on experimental data we know that the dorsal and ventral wavelengths are equal (*λ_d_* = *λ_v_*).

Additionally, and without loss of generality, we can assume that the ventral phase offset can be calculated in relation to the dorsal wave. This simplifies the equation to

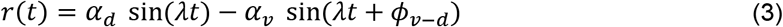

Since the two sine waves have same wavelength, the results of subtracting them will be a new sine wave with the same wavelength, but a different amplitude and phase offset. We can determine these values by treating the dorsal and ventral waveforms as two vectors 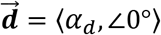 and 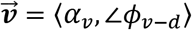 and then calculate the vector 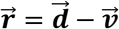 using the trigonometric technique (Hecht, 2017).

Figure 4A shows a conceptual representation of this method where the vectors 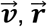 and 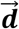 form a scalene triangle. To calculate the amplitude and phase offset of 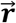, the triangle formed by these vectors can be solved, Using the law of cosines, *α_r_*, the amplitude of 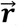, can be calculated with the following equation

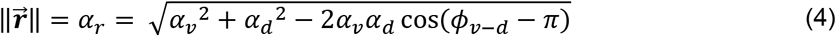

**Figure 4:**
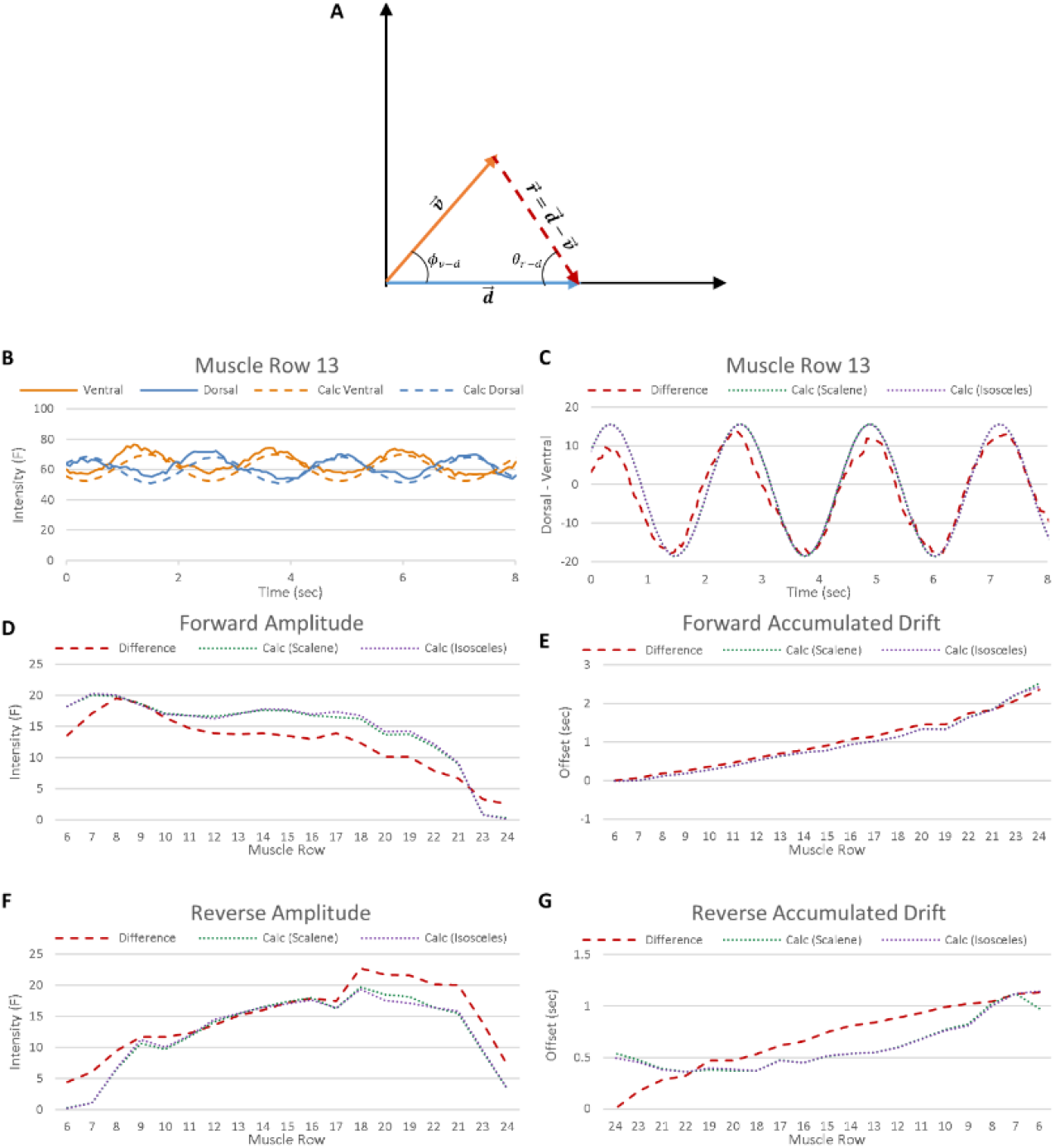
Mathematical analysis of forward and reverse body wave propagation. A) Trigonometric method for determining the resultant phase shift *θ_r–d_* and amplitude 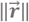 of the muscle activity that results from off-phase dorsal and ventral muscle activation. The vector 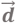 represents dorsal activity, 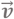 represents ventral activity, *ϕ_v–d_* is the phase difference between 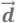 and 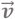, and 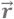 represents the resultant vector. B) Comparison between actual dorsal and ventral signals recorded from muscle row 13 (solid lines) and signals that were generated using the mean, standard deviation, and mean phase shift information computed in this study (dashed lines). The generated signals very closely approximate the recorded activity. C) Comparison of recorded muscle row 13 dorsal-ventral activity and the generated signals that result from applying the scalene and isosceles methods for computing the resultant vector 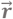. The results of both methods agree well with the recorded activity. D) Comparison of the recorded mean amplitude during forward locomotion and the values computed using the scalene and isosceles methods. E) Comparison of the recorded accumulated drift and calculated accumulated drift during forward locomotion. In agreement with Figure 5B, body wave propagation results from the combined activity of the dorsal and ventral sides. F) Comparison of the recorded mean amplitude during reverse locomotion and the values computed using the scalene and isosceles methods. G) Comparison of the recorded accumulated drift and calculated accumulated drift during reverse locomotion. In agreement with Figure 5J, reverse body wave propagation results from the combined activity of the dorsal and ventral sides.

Similarly, the phase offset between 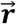 and 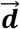, can be computed using the law of sines

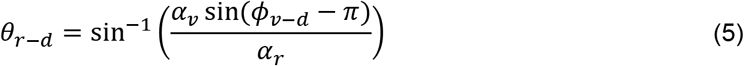

Since the amplitude of the dorsal and ventral sides are nearly equal to one another, *α_d_* ≈ *α_v_*, the triangle formed by 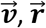, and 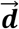 could also be treated as isosceles. Under this assumption, the equation for determining the amplitude of 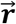 becomes much simpler

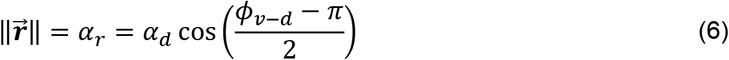

and the equation for determining the phase offset simplifies to

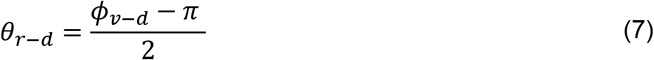

Using standard deviation information from Figures 1B & C, we determined the average peak-to-peak amplitude of the dorsal and ventral sine waves for the muscle cells by using the well know conversion method from Equation 1 (Smith, 1999).

Figure 4B shows a comparison of waveforms recorded for the dorsal and ventral muscles in the 13^th^ muscle row and sine waves that were generated from the computed amplitude, frequency, and phase offset collected during this study. The results are a very close approximation to the actual data. We also generated the sine waves that result from subtracting the ventral from dorsal signals using both the scalene and isosceles trigonometric methods presented in Equations 2–7. Figure 4C shows the comparison between the recorded difference and the computed differences using these two methods.

Again, the results show a very close approximation to the actual data. Finally, Figure 4D–6G show the recorded amplitudes and accumulated drift for muscle rows 6-24 during forward and reverse locomotion and the calculated approximations. The calculated results for forward locomotion amplitude and drift, along with the reverse amplitude, are a very close fit to the actual data. However, the accumulated drift in reverse deviates from recorded values, although the calculated values are still a reasonable approximation.

These results confirm that the overall muscle activation pattern seen in *C. elegans* could result from muscle activation waves that travel down the dorsal and ventral sides at different rates.

## DISCUSSION

### Using high-resolution wavelength switching to measure calcium transients in freely moving worms

There are several advantages and disadvantages to using wavelength switching as the basis for measuring calcium transients in freely moving worms. Unlike other techniques that require projecting multiple images onto the imaging sensor (Butler, et al., 2015), one of the key advantages of performing wavelength switching is the ability to take very high-resolution images of the specimen. Our system, for instance, uses a Zyla-4.2P-CL10 sCMOS camera that can take images at a resolution of up to 4 megapixels (2048 X 2048). If we assume that the length of the worm’s body fills half of the viewable area of the camera, this provides approximately 1024 pixels of resolution along the A-P axis. At such a high resolution, the worm’s body can be partitioned according to the location of the individual muscle cells making it much easier to attribute intensity changes to muscle activity. In addition, by taking large spatial samples of each cell, we can compensate for camera noise by averaging (filtering) the recorded intensities over many pixels. At the resolution and zoom used in collecting our data, for example, each muscle cell is spatially sampled using an average of about 4,500 pixels.

Obtaining images at such high resolution comes at the expense of temporal resolution. Switching between just two wavelengths can cause a significant drop in the frame rate especially if one of the wavelengths comes from a mechanically shuttered light source. On our system, the mechanical shutter on the transmitted brightfield source takes approximately 12 ms to open or close. When combined with the exposure, transfer, processing, compression and storage time, this leads to a sample rate of just 14.2 samples per second.

For measuring calcium transients in muscle cells, this sampling rate is more than adequate. Muscle cells have a calcium transient frequency of approximately 0.45 – 0.48 Hz (Karbowski, et al., 2008). With a sample rate of 14.2 samples per second, our system captures 31 measurements during each wavelength of a muscle cell’s oscillation. Given that the sampling criteria (also known as the Nyquist criteria) only requires a sampling rate that is twice the signal frequency (Shannon, 1949) in order to be identified, our system is more than capable of accurately capturing and characterizing muscle calcium transients. However, if the goal were to capture muscle cell action potentials in freely moving worms using optogenetic techniques with faster kinetics, like the genetically encoded voltage indicator QuasAr (Hashemi, et al., 2019), then the sample rate would need to be increased. According to electrophysiological measurements, a BWM action potential has a duration of 96.3 ± 5.7 ms (Liu, et al., 2011). Therefore, the sample rate would need to be increased to at least 21 samples per second in order to meet the sampling criteria. This could easily be achieved in our system by using binning to reduce image resolution, replacing the mechanically shuttered light source, and/or reducing the exposure time.

### Varying rigidity through muscle activity

It is certainly not unheard of for invertebrates to use their muscles to alter body rigidity. For example, it is believed that lampreys modulate their stiffness using appropriately timed muscle contractions to create rigidity in their tail while swimming. This strategy allows the organism to transfer positive work from anterior muscles through the tail to the surrounding fluid (Tytell, et al., 2010).

Although we do not believe that *C. elegans* uses a similar strategy during crawling, our results suggest that the worm uses its muscles to selectively alter rigidity in areas that are being used to produce the greatest amount of locomotive force. Because our measurements were normalized using maximum, F_4AP_, instead of mean, F_0_, fluorescence intensity, we were able to see that 1) mean intensities are the same in dorsal/ventral contralateral muscle pairs throughout the entire body during both forward and reverse locomotion and 2) there are significantly elevated anterior (posterior) mean intensity levels that linearly decrease to significantly lower posterior (anterior) mean intensity levels during forward (reverse) locomotion. When these results are combined with the findings that GCaMP fluorescence intensity is associated with muscle length (Liu, et al., 2011), that muscle length is associated with cuticle stiffness (Petzold, et al., 2011), and that muscle waves decrease as they travel through the body (Xu, et al., 2018), they support this conclusion.

Our results in no way suggest that previously measured values of the Young’s modulus of *C. elegans* are inaccurate (Park, et al., 2007; Fang-Yen, et al., 2010; Sznitman, et al., 2010). They do, however, highlight that current techniques used to measure this value produce a mean (whole body) value that does not account for varying stiffness due to muscle activity.

### Dorsal and Ventral Neuromechanical Phase Lag

Neuromechanical Phase Lag (NPL) is characterized by an increase in the phase difference between body bending and muscle activation as a locomotion wave travels from source to destination (McMillen, et al., 2008). NPL have been discovered in several organisms including eels, lampreys, and sandfish (McMillen, et al., 2008; Tytell, et al., 2010; Ding, et al., 2013).

Previous theoretical and empirical studies show that *C. elegans* lacks a NPL and instead has a fixed phase offset between bend angle and muscle activation throughout the entire body during forward locomotion (Fang-Yen, et al., 2010; Butler, et al., 2015). Our results agree with these findings and extend them to include reverse locomotion as well. Figure 2B & J show the results of comparing the propagation of the body and combined muscle activation waves for forward and reverse. Because these results are not adjusted for the kinetics of GCaMP, they show an equal accumulated drift over the entire propagation of the wave. These results agree with the raw results recorded by Butler et al. during forward crawling (Butler, et al., 2015). Figure 5 shows kymographs for the bend angle and difference in Ca^++^ intensity between the dorsal and ventral sides for both forward and reverse locomotion. The red lines are overlaid to show zero crossing of each body wave and are copied onto the calcium activation kymographs for reference. Note that the lines are nearly perfectly aligned with the zero crossings of the activation graphs indicating a phase offset of 0 degrees.

**Figure 5:**
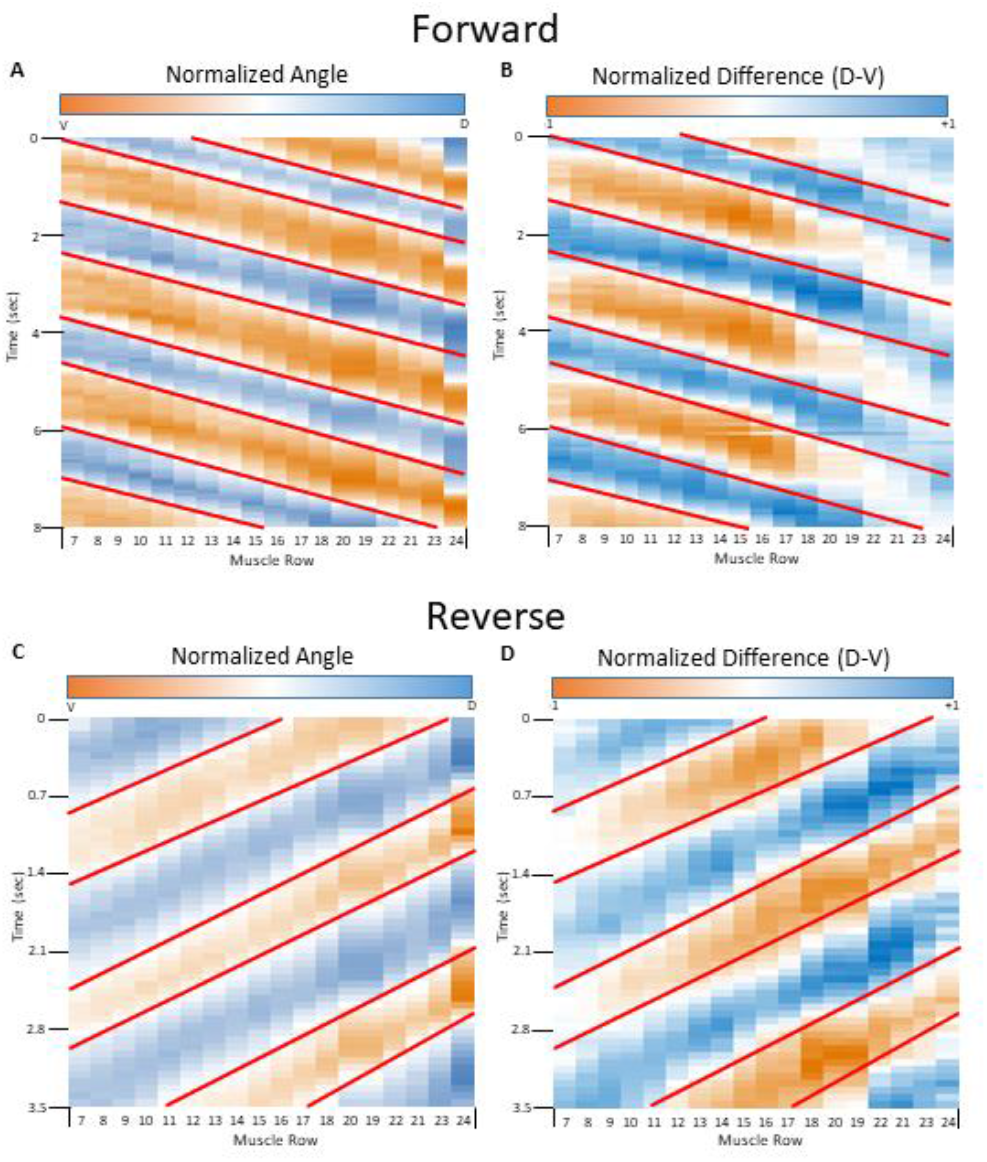
Kymographs of forward and reverse locomotion. A) Sample kymograph of body wave propagation over time during forward locomotion of the same worm and video segment as Figure 7A-I. B) Sample kymograph of the dorsal – ventral Ca++ activation over time during forward locomotion of the same worm and video segment as Figure 7A-I. C) Kymograph of body wave propagation over time during reverse locomotion of the same worm and video segment as Figure 7J-R. D) Kymograph of the dorsal – ventral Ca++ activation over time during reverse locomotion of the same worm and video segment as Figure 7J-R.

Our results, however, also show a previously unreported phenomena where the dorsal and ventral activation waves travel at different rates in relationship to both body bending and each other. Figure 2B, for instance, shows that during forward locomotion, the dorsal activation waves travels 41% faster (1.88s versus 2.66s) than the body bending wave. In addition, the ventral wave travels 57% faster than the dorsal wave (1.18s versus 1.88s). This graph, along with Figure 2J for reverse, show clear NPLs when the two sides of the worm are considered independently of each other.

Given that it is currently believed that the presence of an NPL indicates that wave propagation is primarily driven by one or more CPGs (Tytell, et al., 2010; Denham, et al., 2018), these results lend considerable support to the hypothesis that CPGs play a significant role in the generation and propagation of locomotive waves in *C. elegans*. Our findings suggest that there may be independent dorsal and ventral CPGs that are coordinated with one another during both forward and reverse locomotion.

Several studies have reported finding CPG activity in the ventral nerve cord. Two studies, for example, reported that B-type motor neurons (MN) may function as oscillators to drive forward locomotion (Wen, et al., 2012; Fouad, et al., 2018). Additional studies have reported that AS MNs may form oscillators with DA and DB neurons (Olivares, et al., 2017; Tolstenkov, et al., 2018) and that A-type MNs (particularly DA9) may also be rhythmic in nature (Gao, et al., 2017).

It has also been shown that proprioception plays an important role in regulating the activity of these CPGs (Wen, et al., 2012) and that D and AS MNs work to coordinate the dorsal and ventral sides. For example, AS MNs have been shown to drive dorsal BWMs while simultaneously activating VD MNs that down regulate the activity of ventral BWMs (Tolstenkov, et al., 2018). A recent study of D-type MNs has also shown that impairing the function of the cells leads to decreased coordination, undulation frequencies, and locomotion speed (Deng, et al., 2020).

These impairments could certainly be explained using a model where the two sides of the worm exhibit proprioceptive modulated CPGs that are coordinated with one another. By recognizing that muscle activity in different regions of the body become potentiated and/or attenuated by a lack of proper phase alignment, it is easy to see that removing coordination between the dorsal and ventral sides would result in decreased speed, increases and decreases in bend angles, and altered bend angle frequencies.

## MATERIALS & METHODS

### *C. elegans* Growth Conditions and Strains

*C. elegans* were grown at 20°C on NGM lite plates containing streptomycin and nystatin; seeded with OP50-1 as a bacteria food source (Brenner, 1974). All experiments were completed with young adults, age-synchronized by picking L4 stage animals to fresh food plates 12-24 h before the experiment. Strains used in this study were N2 and HBR4: goels3[myo-3p::SL1::GCamP3.35::SL2::unc54 3’UTR + unc-119(+)] (Schwartz, et al., 2012).

### Single Worm Tracking Protocol

A worm was transferred using a platinum worm pick into an 80μL drop of H20 on a glass microscope slide at room temperature to remove residual bacteria. After 30 seconds, the worm was transferred to a 1.5μL drop of H20 on a modified NGM lite plate (1/20 w/w reduction in peptone). The plate was immediately transferred to the microscope for tracking. The 1.5μL drop of water absorbs into the plate within 2 min allowing for transfer and focusing prior to tracking. Worms were tracked under rapid wavelength switching conditions at room temperature, explained below, and each experiment was concluded after four minutes of recording.

### Image Capture

Images were acquired using a Leica DMi8, X-Cite XLED1 (Lumen Technologies), Zyla-4.2P-CL10 sCMOS camera (Andor), and in-house software (CNAS tracker). During recording, subsequent images were captured under brightfield and GFP (~470 nm) illumination. The synchronization of image capture and wavelength switching was accomplished using an Arduino-based sequencer that triggered the XLED1, brightfield shutter on the DMi8, and Zyla-4.2P. To account for the delay associated with the mechanical shutter on the brightfield light source, a 12 ms delay was introduced between wavelength changes. Images were captured at 2048 x 2048 using a 10x objective and an exposure time of 9.8 ms. Brightfield and fluorescence intensities and aperture settings were unchanged between samples. Frames were stored on a hard drive using a custom video format called Rapid Video Compression (RVC). Each frame was annotated with a time stamp, stage position, and the lighting conditions. Average frame time was 36 ms. Figure 6A – D shows the results of this process for both forward and reverse locomotion. In Figure 6A & C, the brightfield images are shown, while in Figure 6B & D the corresponding GFP images are shown. To emphasis the changing patterns seen in the GFP signal as the worm moves, these images show every third frame pair (191 ms interval) captured by our system.

**Figure 6:**
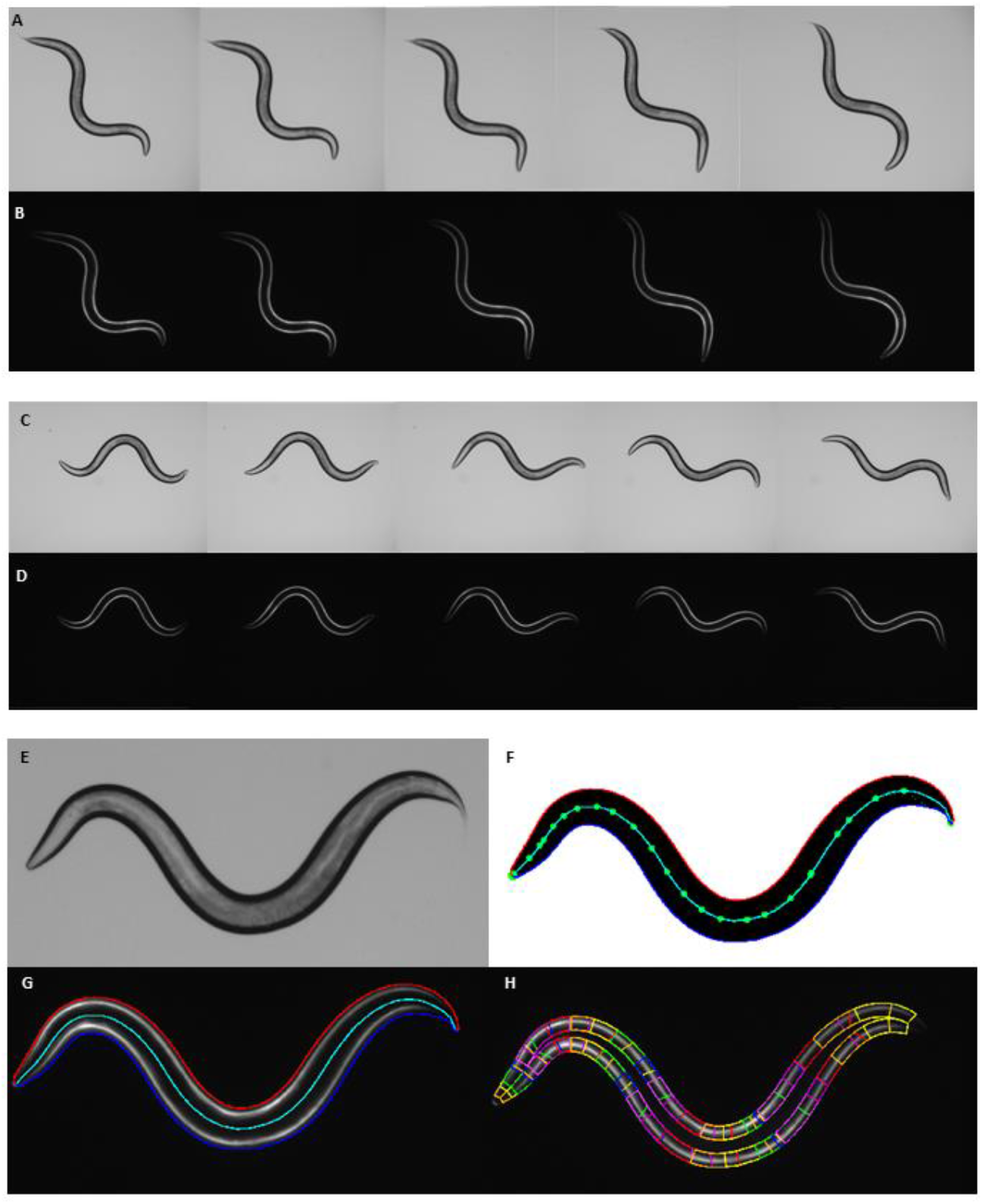
Image processing steps used to quantify bend angles and muscle calcium signals. A) Series of brightfield images of *C. elegans* during forward locomotion. Images are taken at an interval of 191ms. B) Series of corresponding GFP images of *C. elegans* during forward locomotion. Images are taken at an interval of 191ms. C) Series of brightfield images of *C. elegans* during reverse locomotion. Images are taken at an interval of 191ms. D) Series of corresponding GFP images of *C. elegans* during reverse locomotion. Images are taken at an interval of 191ms. E) Bright field image of a freely moving *C. elegans*. F) Bright field image converted to a binary image with the border outline and skeleton. The blue outer border indicates the dorsal side and the red outer border indicates the ventral side. The midline (cyan) is segmented according to the midpoints of the muscle cells (green). Note that muscles 1 and 2 are located directly on top of each other. G) The outline and midline are used as a template for the GFP image. H) Individual outlines for each of the muscle cells are analyzed to determine their Ca++ concentration. To account for cell overlap, a distance weighted average is computed.

### Worm tracking

Worms were tracked using in-house software (CNAS tracker), which controlled the stage on the DMi8. The worm was identified in each brightfield frame by searching from the center of the last bounding region in expanding concentric circles for a seed pixel (a pixel below a given threshold). Once a seed pixel was identified, a threshold-based scan line flood fill algorithm was used to determine a bounding region. An offset between the center of the frame and the center of the bounding region was then calculated and if greater than 50 pixels, the stage was repositioned.

### Video Analysis

Videos were hand partitioned into segments of forward and reverse locomotion using in-house video editing software (RVC editor). Each video segment was then analyzed using in house video analysis software (Worm Analyzer). During video processing, brightfield and GFP frames were analyzed in pairs. Brightfield images (Figure 6E) were converted to binary using a threshold algorithm. Holes were filled in and small objects were removed using a combination of dilation and erosion operations. A seed pixel was then located in a similar manner to the method used for worm tracking.

To determine the shape of the worm’s body, a border following algorithm was used to trace the worm’s outline. The head and tail were then located by finding the maximum point of curvature along the outline, which was identified as the tail, and the second maximum point of curvature, which was identified as the head. The outline was then partitioned into left and right sides based on the head and tail location. These sides were then manually identified as dorsal/ventral using the vulva location as a guide.

The centerline was created by tracing one side of the border and calculating the perpendicular to the curvature at each point. This perpendicular was then intersected with the opposing side of the worm and the center point of this line segment calculated. The center points were added to a data structure that ordered them based on X and Y location and finally, gaps in the centerline were filled in between subsequent centerline points as needed.

The skeleton was constructed by partitioning the centerline into 25-line segments using the head, tail, and midpoints of the BWMs as end points. The midpoint locations of the muscle cells were used to calculate the angles of the body (Figure 6F).

Because the stage can move between the brightfield and GFP images, analysis of the GFP frames began by realigning the outline of the worm with the GFP image. This was accomplished using a template matching algorithm that maximizes the fluorescence signal within the outline (Figure 6G). A bounding region for each muscle was then determined using the outer border, the anterior and posterior extents of the muscle cell, and an inset border. The inset border mirrored the curvature of the outer border and was placed 25 micrometers from the outer border but did not cross the centerline of the worm (Figure 6H).

Once the muscle regions were determined, the average intensity of each muscle cell was calculated using a modified scanline flood fill algorithm. To account for the shape of the muscle cells, weights were assigned to each pixel’s value based on its distance from the center of the muscle cell. This technique placed higher weight on the pixels in the thickest and least overlapping portions of the muscle cell. Relative fluorescence Intensity (RFI) was then calculated by subtracting the average background fluorescence from the weighted fluorescence intensity of the region.

Results of the image analysis were stored in Comma Separated Values (CSV) format files. For each pair of frames, a single line was recorded with the timestamp, X and Y stage location, the 24 angles, and 48 intensities of the muscle cells along the body. Figure 7A-R show traces of the bend angles and muscle cell activation during forward and reverse locomotion for the 3^rd^, 13^th^, and 23^rd^ muscle groups.

**Figure 7:**
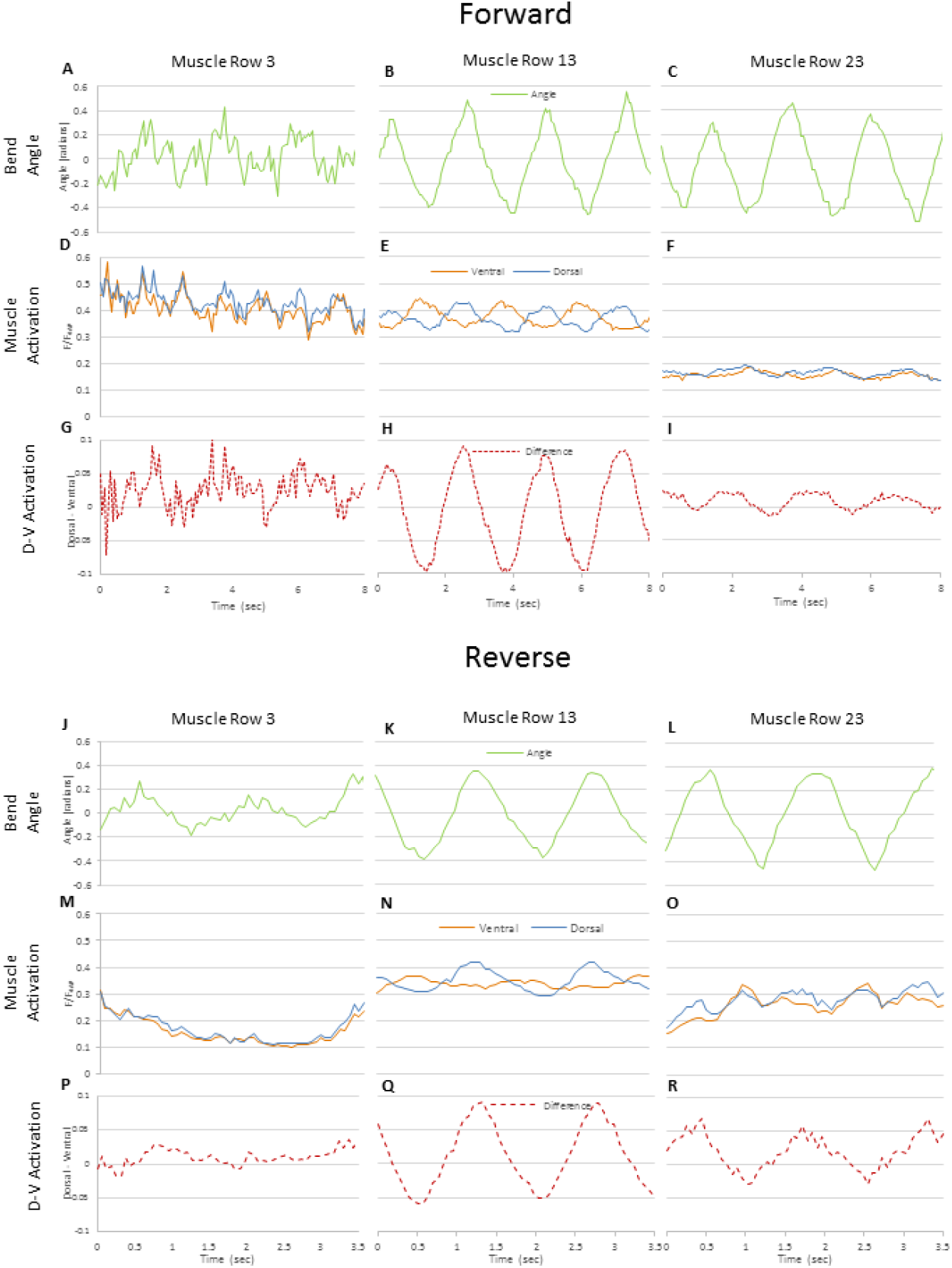
Trace of the bend angles and calcium concentrations for three locations along the AP axis during forward and reverse locomotion. A-C) Forward locomotion bend angle data for the 3^rd^, 13^th^, and 23^rd^ muscle rows, which are located in the head, mid-body and tail, respectively. The head moves rapidly from side to side, while the pattern at the mid-body and tail are very sinusoidal and have nearly the same amplitude. D-F) The normalized GFP intensity of the ventral and dorsal muscles during forward locomotion at the 3^rd^, 13^th^, and 23^rd^ muscle row. At the head, the signals are nearly perfectly in phase and bare a close resemblance to the changes in bend angle. At the mid-body, the pattern becomes clearly anti-phasic with the same amplitude as the head muscles. At the tail, the signals are nearly in phase again, but considerably less intense than the head or mid-body. G-I) The normalized difference between the dorsal and ventral GFP intensities at the 3^rd^, 13^th^, and 23^rd^ muscle rows during forward locomotion. Within the head, the difference in intensity is only weakly correlated with the bend angle. However, at muscle rows 13 and 23, the bend angles and difference in intensities are highly correlated. J-L) Bend angle data for the 3^rd^, 13^th^, and 23^rd^ muscle rows during reverse locomotion. The bend angles are clearly sinusoidal throughout the body. Rapid head movement, however, is suppressed. M-O) Normalized GFP intensities for the 3^rd^, 13^th^, and 23^rd^ muscle rows during reverse locomotion. Head activation is still in phase, but significantly lower than forward locomotion. Tail activation is still in phase, but much higher than during forward locomotion. P-R) The normalized difference between the dorsal/ventral GFP intensities at the 3^rd^, 13^th^, and 23^rd^ muscle rows during reverse locomotion. Head muscles show low levels of sinusoidal activation, while mid-body and tail activity are strongly correlated with body posture.

### 4-Aminopyridine inhibition

To normalize for the expression pattern of myo3p::GCamp3, we conducted experiments to measure the Ca^++^ levels in worms (n=10) that were exposed to the K^+^ channel blocker 4-AP. Exposure to 4-AP causes dramatic and sustained activation of the muscle cells throughout the body because the cells lose the ability to repolarize. This allowed us to record “maximum” Ca^++^ levels for the BWMs (Figure 1A).

One day old adults were incubated in a 60 μL drop of 500 mM 4-Aminopyridine (4-AP) on a glass slide for 6 minutes in a humidified chamber. Following the 6-minute incubation, the worms were transferred directly to a 1.5 μL drop of 500 mM 4-AP on a tracking plate. Once the 500 mM 4-AP absorbed into the plate (this process takes approx. 1 minute), tracking was started and continued for 45 seconds (n=10).

### Data Analysis

The data files produced from the image analysis were batch analyzed using a combination of the Worm Analyzer and Microsoft Excel. During analysis, aggregate amplitude statistics were gathered for each individual body bend angle, body bend speed, muscle cell activation, and the difference between activation of the ventral and dorsal muscle cells in the same muscle row. These statistics included minimum, maximum, mean, and standard deviation (Figures 1A-C and Figures 3A, B, H & I).

The data was analyzed for a second time to generate normalized intensity levels (Figures 1D & E). During this pass, recorded intensity levels were divided by the mean intensity level of the muscle cell recorded during the 4-AP experiment. These values are reported as F/F_4AP_ in the figures.

These files were then processed using custom software written in Java for frequency and correlation information. Frequency information was gathered using a Fast Fourier Transform (FFT) over discrete segments of the input files. Frequency information was aggregated and normalized, so the magnitude reported for each frequency component is the mean magnitude across the segments divided by the maximum mean frequency component.

Correlation analysis was also conducted between ventral, dorsal, activation difference and bend angles (Figure 3C & J). Additionally, autocorrelation was conducted between ventral and dorsal muscles (Figures 2A, C-H and Figures 2I, K-P), and between subsequent muscle rows (Figures 2B & J).

This aggregate data was saved in a text file for further processing in Excel.

### Experimental design and statistical analysis

For HBR4 experiments n=25 worms were tracked. This produced 96 video files of forward locomotion and 111 files of reverse locomotion. In total, 46,226 frame pairs or 3,316.15 seconds of forward locomotion and 6,610 frame pairs or 472.59 seconds of reverse locomotion were analyzed. Average frame rate was ~28 frames per second.

For HBR4 worms exposed to 4-AP, n=10 worms were tracked. Since these worms are paralyzed, the videos did not need to be partitioned between forward and reverse. Therefore, 10 files consisting of 3,570 frame pairs or 303.85 seconds of video were produced. This video averaged ~25 frames per second.

Statistical means and deviations were computed in the standard way. Values were normalized using either the maximum recorded value (for frequency normalization) or the mean value recorded from 4-AP experiments (normalized intensities). FFT analysis was conducted using a window size of 128 samples for forward locomotion resulting in 315 fragments, each about 9 seconds in length. With this window size and an average sample rate of 14.24 samples/second, the maximum detectable frequency is about 7Hz with a frequency resolution of 0.11 Hz.

Because of the short duration of reverse locomotion, the window size of the FFT was reduced to 32 samples. This resulted in 149 fragments that were each approximately 2.2 seconds in length. The average sample rate of this data was 14.6 samples/sec resulting in a maximum detectable frequency of about 7Hz with a frequency resolution of 0.45Hz.

Correlation was computed using the formula for the Pearson correlation coefficient over the data within window (128 for forward and 32 for reverse).

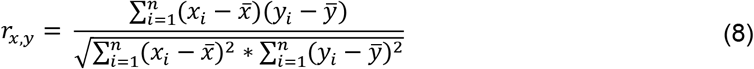

Autocorrelation was conducted by sliding one signal over the other within the window and calculating the correlation coefficient at each offset. The offset was limited to one-half the wavelength in both positive and negative directions. Ventral and dorsal offsets and accumulated drift were calculated using the average offset with the maximum correlation coefficient.

Statistical significance testing was performed using paired, two-tailed t-tests. Data is presented as mean ± SEM using Microsoft Excel unless otherwise noted.

### Code, Software, and Data Accessibility

All code and software developed during this study are available from the corresponding author upon request. All videos and data collected used in this article may also be requested. However, due to the size of the video files they will not be available for download. Under these circumstances, special arrangements can be made.

## ACKNOWLEGEMENTS

Some strains were provided by the CGC, which is funded by NIH Office of Research Infrastructure Programs (P40 OD010440). Special thanks to James Rand for his helpful input and assistance. This work was supported by US Air Force Office of Sponsored Research under grants FA9550-15-1-0060 and FA9550-18-1-0308.

